# Laboratory contamination over time during low-biomass sample analysis

**DOI:** 10.1101/460212

**Authors:** Laura S. Weyrich, Andrew G. Farrer, Raphael Eisenhofer, Luis A. Arriola, Jennifer Young, Caitlin A. Selway, Matilda Handsley-Davis, Christina Adler, James Breen, Alan Cooper

**Affiliations:** Australian Centre for Ancient DNA, University of Adelaide, Adelaide, Australia; ARC Centre of Excellence for Australian Biodiversity and Heritage, University of Adelaide, Adelaide, Australia; Faculty of Dentistry, University of Sydney, Sydney, Australia

## Abstract

Bacteria are not only ubiquitous on earth but can also be incredibly diverse within clean laboratories and reagents. The presence of both living and dead bacteria in laboratory environments and reagents is especially problematic when examining samples with low endogenous content (*e.g.* skin swabs, tissue biopsies, ice, water, degraded forensic samples, or ancient material), where contaminants can outnumber endogenous microorganisms within samples. The contribution of contaminants within high-throughput studies remains poorly understood because of the relatively low number of contaminant surveys. Here, we examined 144 negative control samples (extraction blank and no-template amplification controls) collected in both typical molecular laboratories and an ultraclean ancient DNA laboratory over five years to characterize long-term contaminant diversity. We additionally compared the contaminant content within a homemade silica-based extraction method, commonly used to analyse low-endogenous samples, with a widely used commercial DNA extraction kit. The contaminant taxonomic profile of the ultraclean ancient DNA laboratory was unique compared to the modern molecular biology laboratories, and changed over time according to researchers, month, and season. The commercial kit contained higher microbial diversity and several human-associated taxa in comparison to the homemade silica extraction protocol. We recommend a minimum of two strategies to reduce the impacts of laboratory contaminants within low-biomass metagenomic studies: 1) extraction blank controls should be included and sequenced with every batch of extractions and 2) the contributions of laboratory contamination should be assessed and reported in each high-throughput metagenomic study.

## Main Text

In the new era of culture-independent microbiome research, targeted amplicon or ‘metabarcoding’ approaches are now routinely used to amplify DNA from microbial species across the tree of life. However, these methods lack the ability to select for either specific species or to exclude contaminants [1]. Although these techniques have provided invaluable insight into otherwise cryptic microbial communities, the increased sensitivity and lack of target specificity leaves microbiota studies particularly susceptible to the effects of contamination. Such effects are widespread, as several recent studies have indicated that contaminant microbial DNA can be routinely isolated from laboratory reagents and surfaces [2–4] and that this signal has significantly impacted the interpretation and characterization of microbiota in high-throughput sequencing studies. For example, Salter *et al.*recently demonstrated that bacterial DNA present in laboratory reagents is present in both quality-filtered 16S ribosomal RNA (rRNA) gene and shotgun metagenomic datasets and significantly impacts the interpretation of results [3]. Multiple microbial contaminants have already been identified within the published 1,000 Genomes dataset and other medical genomic studies [4, 5]. Despite these findings, the routine assessment of microbial background contamination is still not required, or fully reported, in microbiota studies.

While the presence of contaminant DNA is widespread, the effects are particularly problematic in low-biomass samples that contain very little endogenous DNA [6] (*e.g.* preterm infant swabs, tissue samples, such as placenta, tumour biopsies, or breast tissue, and some environmental samples, such as ice or calcite). In low-biomass samples, a small contaminant signal from laboratory reagents can easily overpower the intrinsic signal from the sample. This is similarly an issue in current palaeomicrobiology studies that examine ancient, degraded microbiota, such as mummified human tissue, preserved faeces (coprolites), or calcified dental plaque (calculus) [6–8]. In ancient samples, the amount of endogenous DNA attributed to the original source can be extremely low (*e.g.*<0.05% of the total DNA in the sample) and is damaged, fragmented, and intermixed with longer, higher-quality modern DNA fragments from contaminant species [9]. Therefore, monitoring and understanding the contributions of contaminant DNA, especially in low-biomass or ancient samples, is critical to ensure that reported results are only based on the endogenous DNA.

Microbial contaminant DNA (*i.e.* background or exogenous DNA) is a mixture of DNA from both environmental and laboratory sources, with the former including factors such as soil at a burial site, air within the sampling facility, and microorganisms from people touching the sample, while the latter involves reagents, glassware, labware, and surfaces [7]. Environmental contamination in low-biomass samples may be difficult to control or monitor, but the laboratory contaminants can be monitored by including extraction blank (EBC) and no-template amplification (NTC) controls and assessed using bioinformatics tools (*e.g.* SourceTracker [10]). An EBC is an empty tube introduced during the extraction steps to collect DNA from the laboratory environment and the reagents throughout processing [11]. Similarly, a NTC is simply an amplification reaction that lacks the addition of DNA from biological samples. These controls should be amplified and sequenced along with other samples and are critical steps to identify and exclude contaminant taxa from downstream analyses, reducing noise and ensuring any results are based solely on endogenous DNA [12]. Despite this, there are surprisingly few published resources describing contaminant taxa found in extraction blank or no-template controls [3, 13, 14].

In this study, we used 16S rRNA metabarcoding to characterise the contaminant diversity in 144 EBCs and NTCs using laboratory techniques specifically designed for low-biomass material. We also explored differences in microbial contamination within two different types of laboratory facilities: a state-of-the-art, purpose-built ancient DNA clean laboratory over the course of five years, and three typical modern molecular biology laboratories over one year. Lastly, we investigated differences between a common commercial DNA extraction kit and a homemade DNA extraction method typically applied in the ancient DNA field. Overall, this study is designed to assess contaminant profiles over time and identify more potential contaminant sequences in both high- and low-biomass research.

## Materials and Methods

### Sample collection

Four different types of sample were used: ancient dental calculus (calcified dental plaque), modern dental calculus, EBCs, and NTCs. Dental calculus samples were obtained from ancient and modern humans as described by Adler *et al.* [11]. A single EBC was included in each batch of extractions by treating an empty tube as if it was a biological sample throughout the DNA extraction and library preparation process. Similarly, NTC samples were created during the 16S rRNA library amplification stage by processing tubes without adding any known template DNA. Both EBCs and NTCs were subsequently included through to DNA sequencing a ratio of one control sample for every ten biological samples.

### Description of laboratory facilities

DNA extraction occurred in two different types of laboratory facilities: a purpose-built, ultra-clean ancient DNA laboratory (ancient lab) and three typical modern molecular biology laboratories (modern labs). The ancient lab is physically remote from the university campus in a building with no other molecular biology laboratories and contains a HEPA-filtered, positive pressure air system to remove DNA and bacteria from external sources. The HEPA filter function is checked annually and changed every ten years. The surface and floors within the laboratory are cleaned weekly with a 5% bleach (NaClO) solution and are illuminated with ceiling mounted UV lights for 30 minutes each night. UV light bulbs are changed annually. Users entering the ancient lab are required to have showered, wear freshly laundered clothing, avoid the university campus prior to entry, and cannot bring personal equipment (*e.g.* phones, writing equipment, and bags) into the facility. Standard personal laboratory wear includes disposable full-body suits, surgical facemasks, plastic see-through visors, and three layers of gloves to allow frequent changing without skin exposure (including one inner elbow-length pair of surgical gloves). All liquid reagents within the ancient lab are certified DNA-free, and the outer surface of all plastic ware and reagent bottles are decontaminated prior to entering the laboratory (cleaned with 5% bleach and treated with UV (2x, 40W, 254nm UV tubes at a distance of 10cm for 10 minutes) within a UV oven (Ultra Violet Products). All DNA extractions and amplification preparations are performed in a room separate to sample preparation and are completed in still-air cabinets that are cleaned with bleach and UV treated for 30 minutes (3x, 15w, 253.7nm tube lamps; AURA PCR) prior to beginning any work. In addition, ancient samples from different sources (*e.g.* soil, plants, and other animals) are processed in separate, dedicated rooms to minimise cross-contamination. In contrast, the modern laboratories are located over 2 km away from the ancient lab at the University of Adelaide (n=2) and the University of Sydney (n=1). All three modern labs are typical of most molecular biology laboratories and are not routinely decontaminated and contain users that routinely use latex gloves but are not required to wear body suits or masks. DNA extracted within the modern labs comes from a wide range of sources (*e.g.* humans, mammals, and environmental samples), although microbiome extractions were only performed on days when no other material was being extracted. In all facilities, DNA was extracted and prepared for amplification in still-air cabinets that are cleaned before and after each use with 5% bleach.

#### DNA extractions

Several specialized DNA extraction protocols have been developed within ancient DNA studies to remove environmental contamination and enhance the recovery of the endogenous DNA. The extraction method selected for this study has previously been described for work on ancient dental calculus [12]. Each ancient sample was first decontaminated using a published protocol [11], while modern samples were not decontaminated. The decontamination procedure included exposure to UV radiation for 15 minutes on each side of the sample, submersion of the sample in 5% bleach for 5 minutes, followed by submersion in 90% ethanol for 3 minutes to remove any residual bleach, and 5 minutes of drying. Decontaminated ancient calculus was then wrapped in aluminium foil and pulverized into power with a steel hammer and placed into a sterile 2mL tube. The EBCs were empty tubes exposed to air for 30 seconds in the same room during sample decontamination and were included in the extraction process as if they contained a sample.

Following decontamination, DNA was extracted using the QG-based method previously described for the extraction of ancient microbiome material [12] (referred to as ‘QG’). All reagents for the QG extraction method were prepared in a ‘sample-free’ room in the ancient DNA facility, and all reagents were aliquoted immediately upon opening and frozen until further use to avoid cross contamination. Where possible, certified ‘DNA-free’ reagents and lab ware were purchased (*e.g.* water and plastic tubes). All other reagents were opened solely within a sterilized hood within the ancient DNA facility. All chemicals were prepared for the extraction with previously unopened DNA and RNA-free certified water (Ultrapure water; Invitrogen). Briefly, 1.8 mL of 0.5 M ethylenediaminetetraacetic acid (EDTA; Life Tech), 100 µL of 10% sodium dodecyl sulphate (SDS; Life Tech), and 20 µL of 20 mg/mL proteinase K (proK; Life Tech) were added to each sample, and the mixture was rotated at 55°C overnight to decalcify the sample. Released DNA was then purified by adding silica (silicon dioxide; Sigma Aldrich) and 3 mL of binding buffer (*e.g.* QG buffer; Qiagen; modified to contain 5.0M GuSCN; 18.1mM Tris-HCl; 25mM NaCl; 1.3% Triton X-100) [15]. The silica was pelleted, washed twice in 80% ethanol, dried, and resuspended in 100 µL of TLE buffer (10mM Tris, 1mM EDTA, pH 8) twice to elute the DNA, which was then stored at −20°C until amplification. All chemicals were prepared for the extraction with previously unopened DNA and RNA-free certified water (Ultrapure water; Invitrogen). For QG extractions performed in the modern laboratories, unopened aliquots of DNA extraction reagents were transported to the modern laboratory, and the modern samples were extracted following the ancient DNA approach described above.

In contrast to ancient DNA extractions, many modern microbiome studies decrease cost and time by using commercial DNA extraction kits to isolate DNA. In order to compare the nature and extent of contaminant DNA in the ancient method to a typical commercial microbiome DNA extraction kit, we analysed an additional set of EBCs created during extractions using a PowerBiofilm^®^ DNA Isolation Kit (MOBIO) from concurrent oral microbiome research conducted in the same modern labs (referred to as ‘kit’ EBCs).

#### Library Preparation

To minimise additional variables, a simple 16S rRNA amplicon sequencing approach was used in this study to compare the different sample types. Briefly, the V4 region of the bacterial 16S rRNA encoding gene was targeted for amplification using degenerate Illumina fusion primers, as previously described [1]: forward primer 515F (AATGATACGGCGACCACCGAGA TCTACACTATGGTAATTGTGTGCCA GCMGCCGCGGTAA) and barcoded reverse primer 806R (CAAGCAGAAGA CGGCATACGAGATnnnnnnnnnnnnAGTCAGTCAGCCGGACTACHVGGGTW TCTAAT) [1]. The string of n’s in the reverse primer refers to the unique 12 bp barcode used for each sample. Primers were resuspended in TLE buffer within the ancient facility and distributed to the modern laboratory. In both facilities, all PCR amplification reactions were prepared using ultraclean reagents with strict ancient DNA protocols [9]. Each PCR reaction contained 17.25 µL DNA-free water (Ultrapure water; Invitrogen), 2.5 µL 10X reaction buffer (20 mM Tris-HCl, 10 mM (NH_4_)_2_SO_4_, 10 mM KCl, 2 mM MgSO_4_, 0.1% Triton^®^ X-100, pH 8.8@25°C; ThermoPol Buffer; New England Biolabs;), 0.25 uL Taq polymerase (Platinum Taq DNA Polymerase High Fidelity; Thermo Fisher Scientific), 1.0 µL MgCl_2_ (Thermo Fisher Scientific), 1.0 µL of each primer at 10 uM (IDT), and 2.0 µL of genomic DNA; each reaction was performed in triplicate. 16S rRNA amplification occurred under the following conditions: 95°C for 5 min; 37 cycles of 95°C for 0.5 min, 55°C for 0.5 min, 75°C for 1 min; and 75°C for 10 min. NTC reactions were also included in triplicate. PCR products were quantified (QuBit; Thermo Fisher Scientific) and pooled in batches of 30 samples at equal nanomolar concentrations prior to purification (Ampure; New England Biolabs). Each pool of purified PCR products was quantified (TapeStation; Agilent) before being combined into a single library. All amplicons were sequenced using the Illumina MiSeq 2×150 bp (300 cycle) kit.

#### Bioinformatics Analysis

After sequencing, fastq files for the forward and reverse reads were created using the Illumina CASAVA pipeline (version 1.8.2). Overlapping forward and reverse reads were joined (based on a maximum of 5% nucleotide difference over a minimum 5bp overlap) using BBmerge (sourceforge.net/projects/bbmap/). Only successfully merged sequences were used in downstream analyses. The resulting fastq file was then imported into QIIME (MacQIIME v1.8.0), a bioinformatics pipeline- based software for the analysis of metagenomic data [16]. All further analysis of the amplicon datasets was conducted within the QIIME package. Libraries were demultiplexed using a Phred base quality threshold of less than or equal to 20, with no errors allowed in the barcodes. Operational taxonomic units (OTUs) were determined by clustering sequences at 97% similarity using UClust [17], and representative sequences (*i.e.*cluster seed) were selected for each cluster. By default, clusters with fewer than five sequences were eliminated from the analysis to reduce noise and spurious findings. Lastly, 16S rRNA gene sequences were given taxonomic assignments using the Greengenes 13_8 database if the sequence was at least 80% similar [18, 19]. Taxonomic diversity measurements (alpha- and beta-diversity) and statistical analyses were performed and visualized in QIIME. Samples were rarefied to a minimum of 150 sequences (Figure 2) and a maximum of 1,000 sequences for diversity analyses, as many controls contained low sequence counts. Statistical differences between groups were identified using a PERMANOVA test for beta diversity (adonis), nonparametric t-test for alpha diversity (Monte Carlo), or Kruskal-Wallis and G-tests for detection of specific taxa associated with different treatments.

### Results

#### Low bacterial diversity is routinely obtained from laboratory extraction controls

The EBCs and NTCs were sequenced alongside the ancient and modern biological samples; all sample types were pooled together at equimolar concentrations. Despite the equimolar pooling, we routinely obtained fewer reads from control samples (EBCs and NTCs) compared to the dental calculus samples, likely due to poor amplification of control samples, the quantification of poor DNA libraries, and clean-up strategy employed. Compared to the ancient and modern calculus samples, 6.4-fold fewer reads on average were obtained from EBCs, and 7.6- fold fewer were obtained from NTCs (Figure 1A). As well as containing fewer reads overall, the control samples contained fewer taxa that could be identified than the biological samples. In the ancient laboratory, 719 total OTUs were observed in ancient biological samples (calculus), while only 415 were identified in the EBCs and 228 in NTCs (Figure 1B). In the modern laboratories, 286 total OTUs were described in the modern calculus samples, versus 208 in the EBCs and 102 in the NTCs. The OTU diversity that appears within the EBCs is similar to the differences in diversity observed between modern and ancient biological specimens, potentially reflecting minor cross contamination during DNA extraction. Across different extraction methods, the EBCs for the commercial extraction kit contained 261 OTUs, around 25% more than the in-house method conducted in the modern laboratory. Overall, the laboratory controls were largely dominated by a single phylum, Proteobacteria (Figure 2), and alpha diversity was significantly lower than in the biological samples extracted within the same laboratory (Monte Carlo; p=<0.0001 and T=>11.0 in all comparisons between any group of controls and all biological samples). While the diversity within laboratory controls was considerably lower than the biological samples, these results demonstrate the extent of background microbial contamination even within an ultra-clean laboratory with ‘DNA-free’ reagents, and clearly highlight the need to routinely monitor and report background contamination within all research facilities.

**Figure 1:**
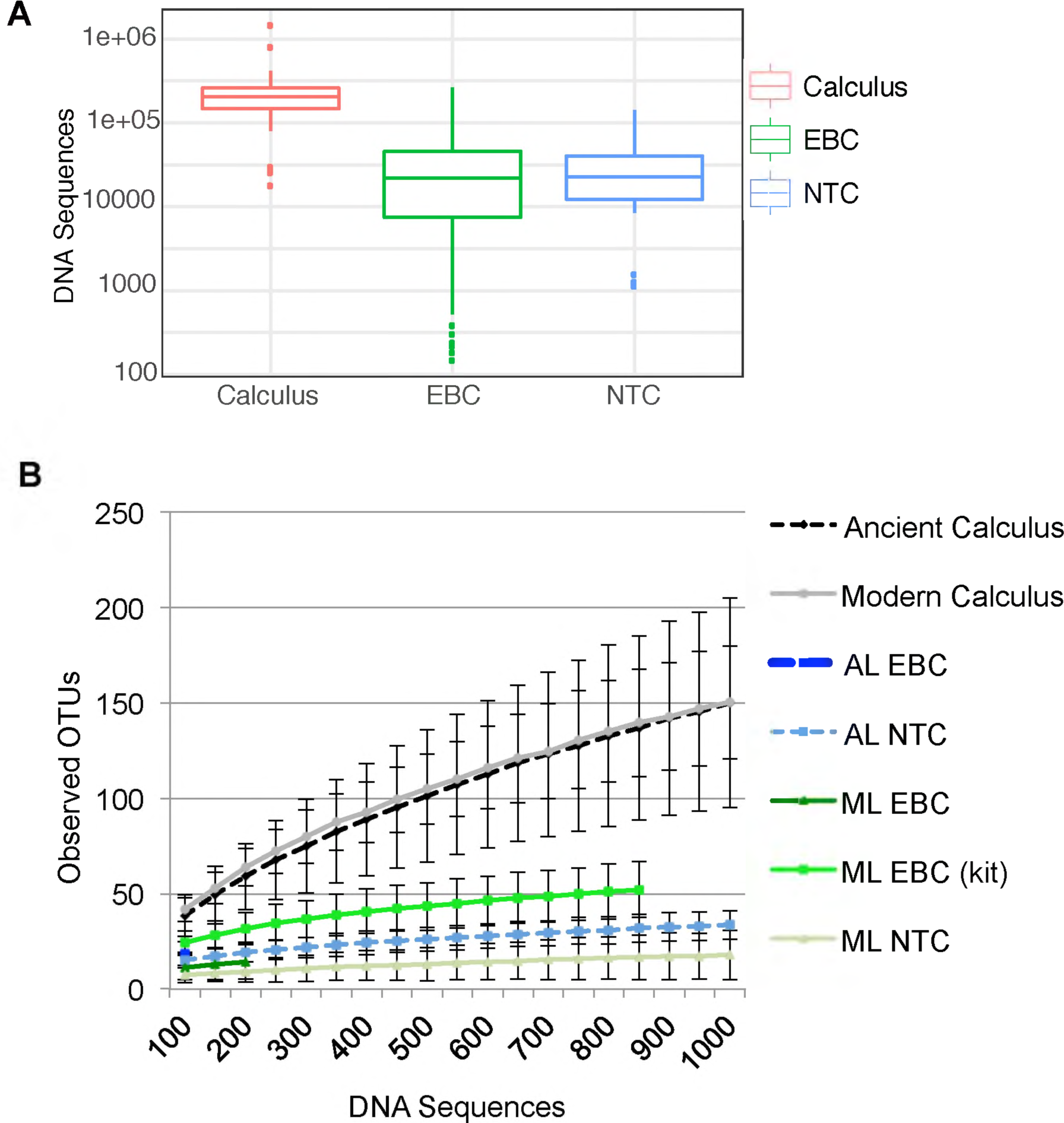
Lower diversity is observed in EBC and NTC samples. The number of sequenced reads from samples that were all pooled at equimolar concentrations is displayed on a box and whisker plot. (B) The alpha diversity of each type sample (*i.e.* the within sample diversity) was calculated using observed species metric in QIIME for rarefied 16S rRNA data. Each sample was rarefied up to 10,000 sequences in 500 sequence intervals; the standard error at each subsampling event is displayed. Calculus samples are shown in blue, while control samples (extraction blank controls (EBCs) and no-template controls (NTCs)) from the ancient laboratory (AL) and the modern laboratory (ML) in red and green, respectively.

**Figure 2:**
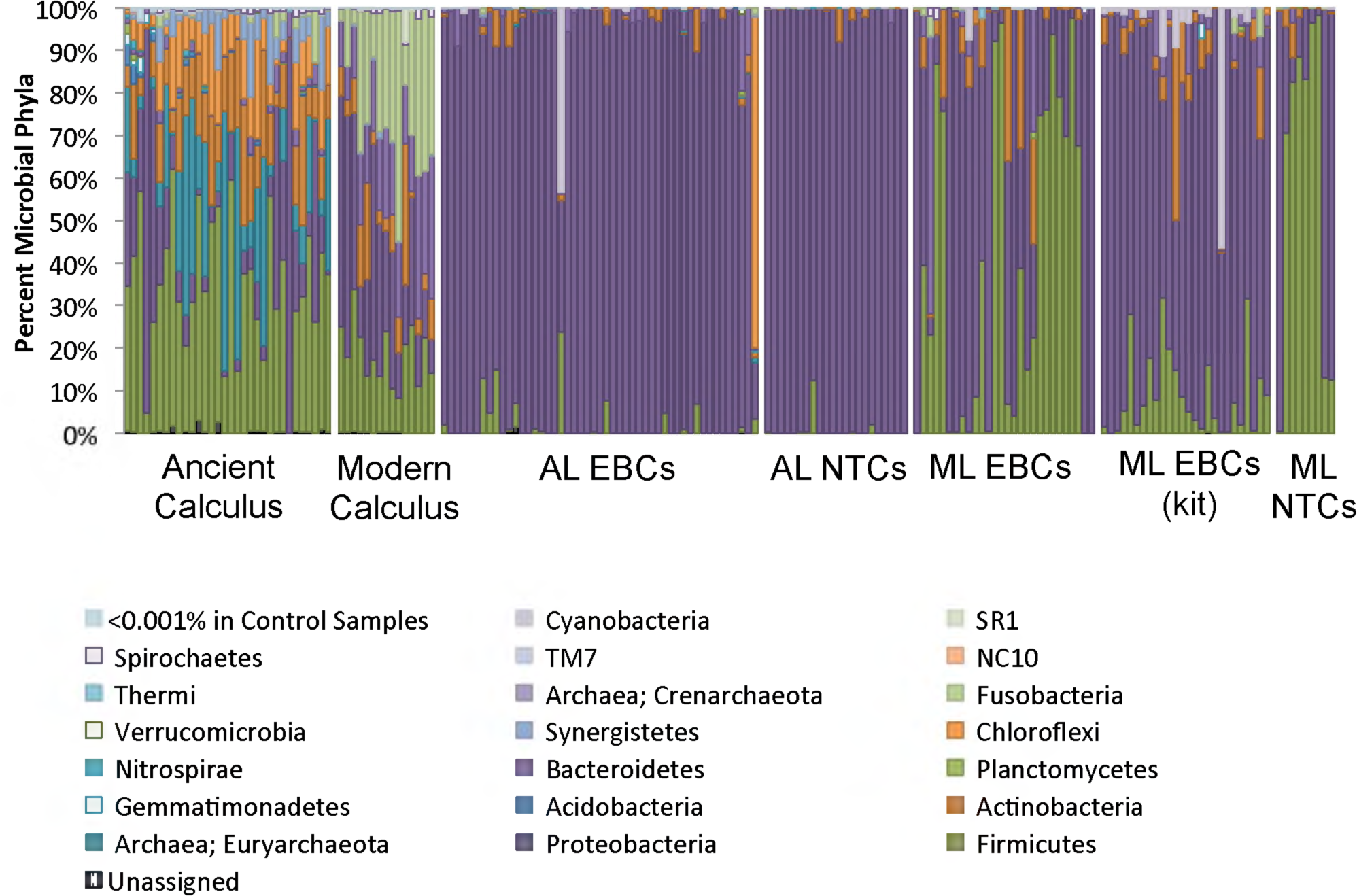
Microbial phyla within controls are distinct from biological samples. The proportion of different microbial phyla are shown for a wide-array of modern and ancient calculus samples and controls samples (EBCs and NTCs) from both laboratory facilities (modern lab (ML) and ancient lab (AL)) and two different extraction methods: the method employed in ancient DNA research and a commercially available DNA extraction kit (kit). Rare phyla were collapsed if the represented less than 0.001% of the total phyla observed.

#### Extraction blank controls detect >50% more contaminant taxa than no-template controls

Many studies, including some in palaeomicrobiological research, have simply reported failed EBC and NTC amplification reactions (often via simple visual comparison on an agarose gel) as a means to determine that their samples are free from contamination [21, 22]. This approach is clearly inadequate, and importantly, also fails to appreciate the extent of contamination introduced during the extraction process, even though this issue is well described in the literature [14, 23, 24]. In our comparisons, EBCs were taxonomically far more diverse than NTCs (Figure 1B) and contained more microbial genera (415 versus 228 genera in the ancient lab, and 208 versus 102 genera in the modern labs). This pattern suggests that if just NTCs were used to monitor the presence of laboratory contamination, at least 53% of the total laboratory contamination may go undetected. These results highlight the need for the standard reporting of both EBCs and NTCs in both modern and ancient metagenomics research.

We examined the impact of overall laboratory contamination on ancient samples by bioinformatically filtering (removing) all contaminant OTUs from ancient dental calculus samples. For the ancient samples prepared with the specialised facility, an average 92.5% of the sequence reads were contaminants, but importantly, only accounted for 28% of the genera identified within these samples. This indicates that endogenous signal can be identified even in low-endogenous samples once contaminant taxa are removed.

#### Extraction blank and no-template controls reflect laboratory environment

Previous studies have detected differences in the contaminants present in different laboratory facilities [3]. In our study, the laboratory environments explained more of the taxonomic diversity observed in the EBCs and NTCs than the extraction or amplification methods used to generate them (Figure 3). For example, Proteobacteria dominated the EBCs and NTCs from the ancient laboratory, while Firmicutes were more dominant in EBC and NTC controls from the modern laboratories. In fact, different types of controls (*i.e.* EBC or NTC) from the same laboratory clustered with others of the same sample type in a Principle Coordinates Analysis (PCoA) of unweighted UniFrac values (p=<0.001, R^2^=0.083; Figure 3A), despite large variation and significant differences in each lab (Figure 1B). Despite the sample type (*e.g.*EBC or NTC) driving the majority of the signal, taxa distinguishing each laboratory could also be detected, with specific *Paenibacillus* taxa only found in the modern laboratories, while the ancient laboratory contained both bacterial (*Comamonas*, *Pseudomonas*, *Acinetobacter*, *Enterobacter*) and archaeal (*Methanobrevibacter*) taxa that were not observed in the modern labs. In addition, several bacterial taxa were identified in both lab types, but were significantly increased in one location. The ancient laboratory contained significantly higher levels of certain *Acinetobacter*, *Comamonas*, and *Pseudomonas* taxa compared to the modern laboratories (Kruskal-Wallis; Bonferroni-corrected p=<0.05), while Erythrobacteraceae and *Staphylococcus* taxa were increased in abundance in the modern laboratories. With the exception of the *Staphylococcus* taxa, each of these taxa had been previously identified in laboratory reagents [3]. This suggests that some contaminant taxa are relatively universal across laboratories and are therefore either introduced in the manufacturing of laboratory reagents and labware or have a fundamental niche in low-nutrient, laboratory environments.

**Figure 3:**
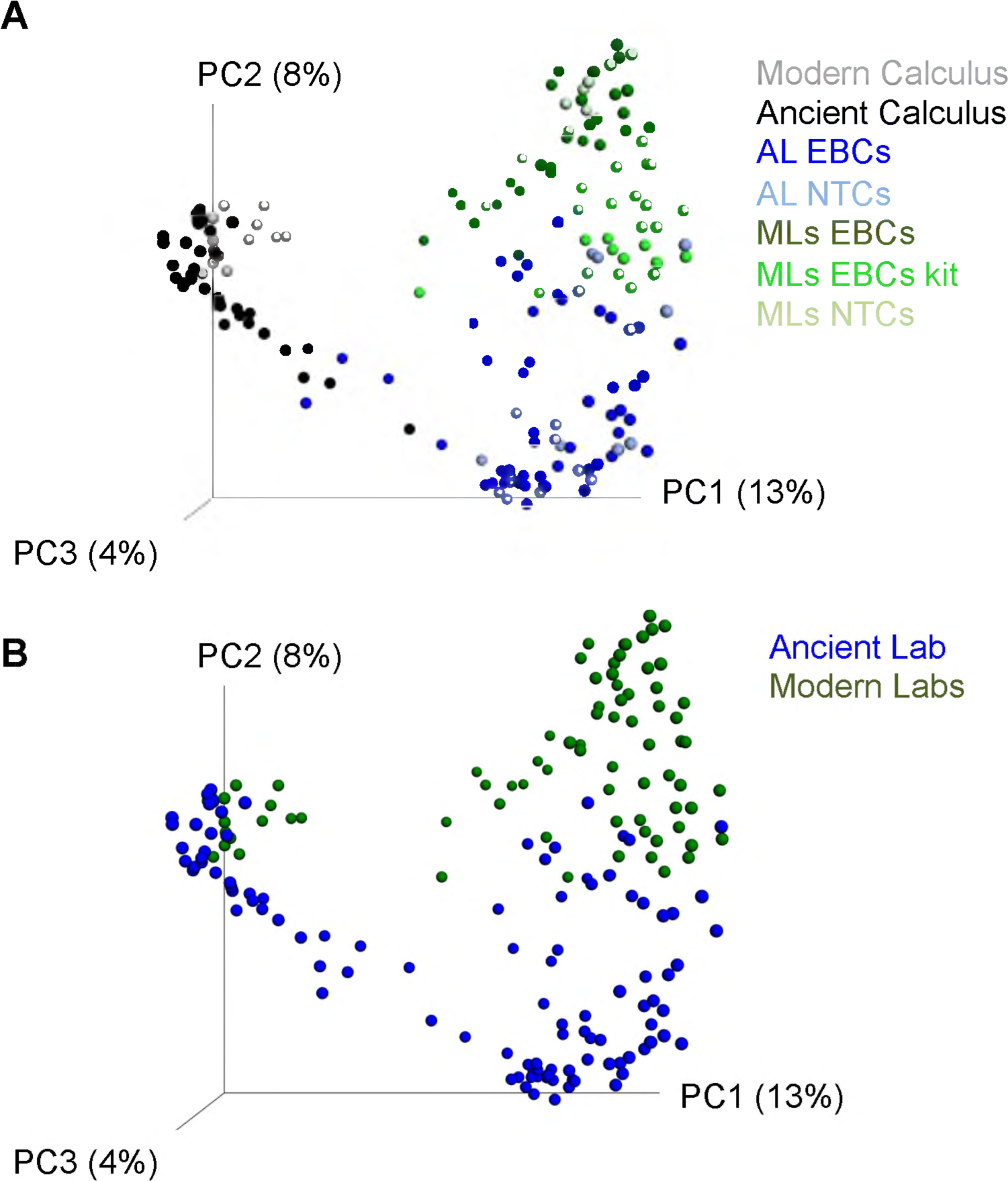
PCoA plots of control samples highlight differences in method and laboratory. PCoA plots of unweighted UniFrac values were plotted in QIIME to compare beta diversity differences (between samples differences) in all samples (A) or in different laboratories (B). The different laboratory facilities are represented by ML (modern lab) and AL (ancient lab), and the two control types are represented by EBC (extraction blank control) or no-template control (NTC).

We next examined the genera that were likely to be in the reagents themselves, rather than the laboratories, by looking for shared taxa within the EBCs generated during extractions in both the ancient lab and modern labs. Of the 69 dominant genera (*i.e.* observed at >0.1%), 17 were present in the reagents used in the in-house QG DNA extraction process used in both types of facility. These taxa included *Cloacibacterium*, *Flavobacterium*, *Paenibacillus*, *Novosphingobium*, *Sphingomonas*, *Limnohabitans*, *Tepidomonas*, *Cupriavidus*, *Ralstonia*, *Acinetobacter*, *Enhydrobacter*, *Pseudomonas*, and *Stenotrophomonas*, and four unidentified genera within Comamonadaceae, Erythrobacteraceae, Enterobacteriaceae, and Pseudomonadaceae (Table 1). Within the ancient laboratory EBCs, the 26 most dominant genera included *Acinetobacter* (39%), followed by three genera within the Comamonadaceae family (totalling 11.3%), *Pseudomonas* (8%), *Novosphingobium* (1.5%), *Ralstonia* (1%), *Cloacibacterium* (1%), and others (Table 1). In the EBCs from the modern laboratories, *Paenibacillus* was the most prevalent of the 43 dominant genera (46%), while two Erythrobacteraceae (16.5%), Comamonadaceae (6.1%), *Cloacibacterium* (3.9%), *Corynebacterium* (2.5%), *Enterococccus* (2.5%), *Staphylococcus* (2.2%), *Enhydrobacter* (1.8%), Microbacteriaceae (1.7%), a Pseudomonadaceae (1.4%), *Ralstonia* (1.3%), and *N09* (1.2%) taxa were the next most prevalent within the reagents (Table 1). Although the same extraction method and reagents were used, only three of the dominant taxa (*i.e.*identified at >1% prevalence) were the same within both laboratories (Comamonadaceae, *Cloacibacterium*, Pseudomonadaceae), highlighting the heterogeneity of taxa identified with EBCs. While many of these taxa have been previously identified as laboratory contaminants, the diversity within the modern laboratories also includes some human-associated taxa that have been cultured from the oral cavity, gut, and skin (*e.g. Corynebacterium*, *Enterococcus*, and *Staphylococcus*, respectively). This suggests that the additional precautionary measures used within the ancient laboratory help reduce the introduction of human-associated microorganisms in metagenomic data sets.

**Table 1:**
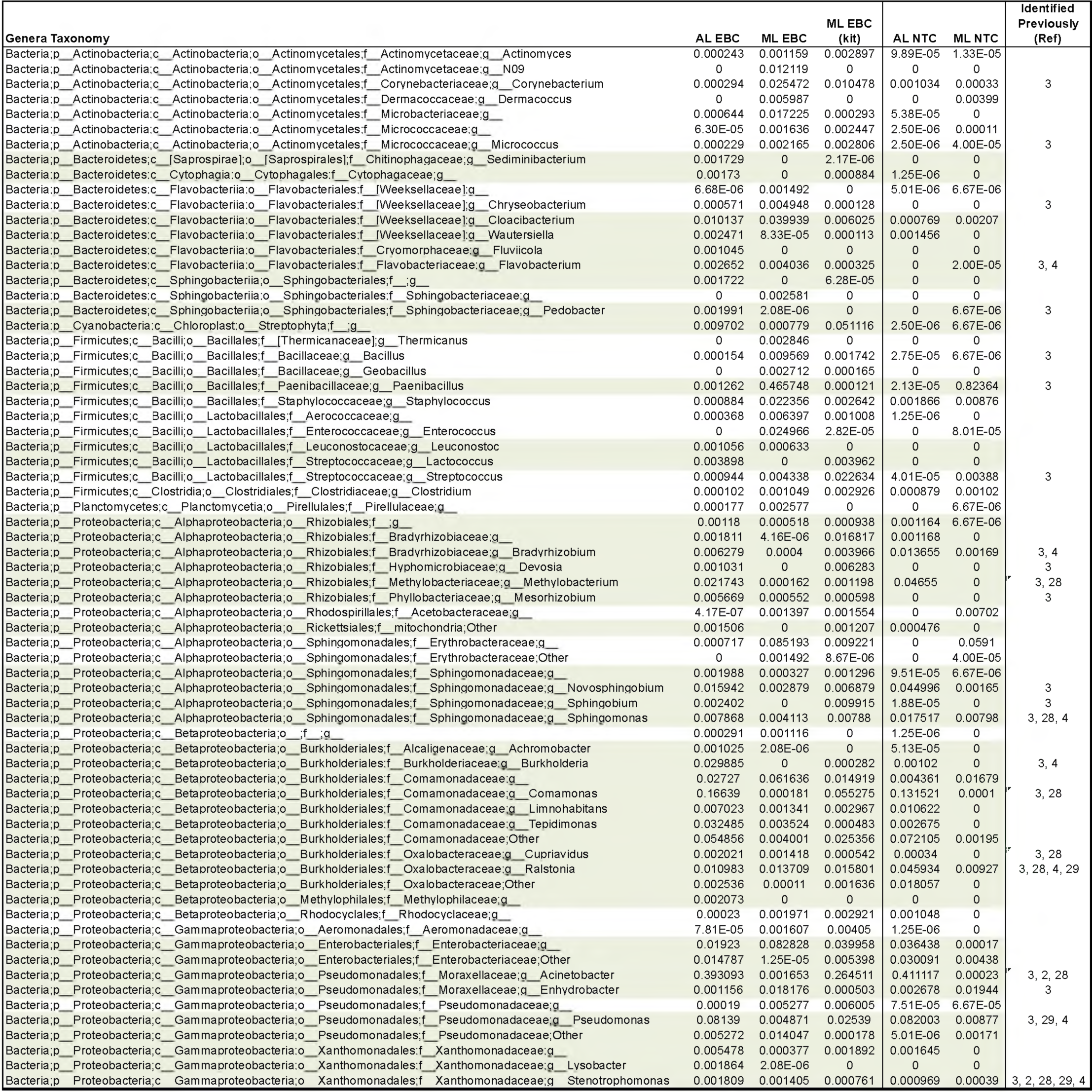
Dominant contaminant genera are largely unique within each laboratory. The 69 genera that dominated EBC control samples are displayed for all sample types and include the proportion identified in each sample type. Genera were identified if dominant if they were found to be above 0.01% of the total genera identified within each laboratory. Taxa highlighted in green represent genera that dominated EBCs in the ancient laboratory, while unhighlighted are those from the modern EBC samples. If the genera were identified in previous studies that examined contamination, the reference number is shown in the right hand column.

#### DNA extraction kits contain microbiota indicative of the human mouth

We compared the diversity of taxa present within EBCs from the widely used ancient DNA extraction method and the commercial PowerBiofilm^®^ DNA Isolation Kit, used in the same modern laboratory. While the latter kit has been shown to have the lowest bacterial background contamination of standard microbiome kits [3], microbial diversity within the kit EBCs was significantly higher than the in-house QG method (Figure 1B), suggesting that kit-based DNA extractions are more prone to background contamination. On a PCoA plot constructed using unweighted UniFrac distances, the kit EBCs clustered away from the QG EBCs and NTCs, including those processed in the same laboratory (adonis; p=<0.001, R^2^=0.04; Figure 4A), demonstrating that a unique microbial community profile originates from the kit. This profile was not solely dominated by Firmicutes, like the other control samples from the modern lab, but contained taxa from several unique phyla (Acidobacteria, Gemmatimonadetes, and Verrucomicrobia). These unique phyla included 15 distinct taxa that were also not observed in the extractions using the ancient DNA extraction method, including *Alicyclobacillus*(n=9), *Halomonas*, *Pseudonocardia*, *Vogesella*, *Allobaculum* (n=2), and *Akkermansia* taxa (Kruskal-Wallis; p=<0.05; Table 2). Several of these taxa are known to be resistant to sterilization treatments, including pasteurization [25]. In addition, several OTUs were more likely to be found in higher abundances in the kit EBCs than any other control samples (G-test; p=<0.05) and include specific *Bradyrhizobiaceae*, *Neisseria*, *Corynebacterium*, *Fusobacterium*, *Streptococcus*, *Micrococcus*, and *Halomonas* taxa. While *Bradyrhizobium* and *Micrococcus* have previously been identified as laboratory contaminants [3, 4], the remaining taxa are commonly found in the human mouth. Concerningly, many of these human oral taxa have been previously reported from low-biomass samples, such as placenta and tumor tissue, which were examined without EBCs [22, 26]. This suggests that DNA extraction kits used in modern molecular biology laboratories may be contributing unique microbial signals in addition to those generated within the laboratory environment.

**Figure 4:**
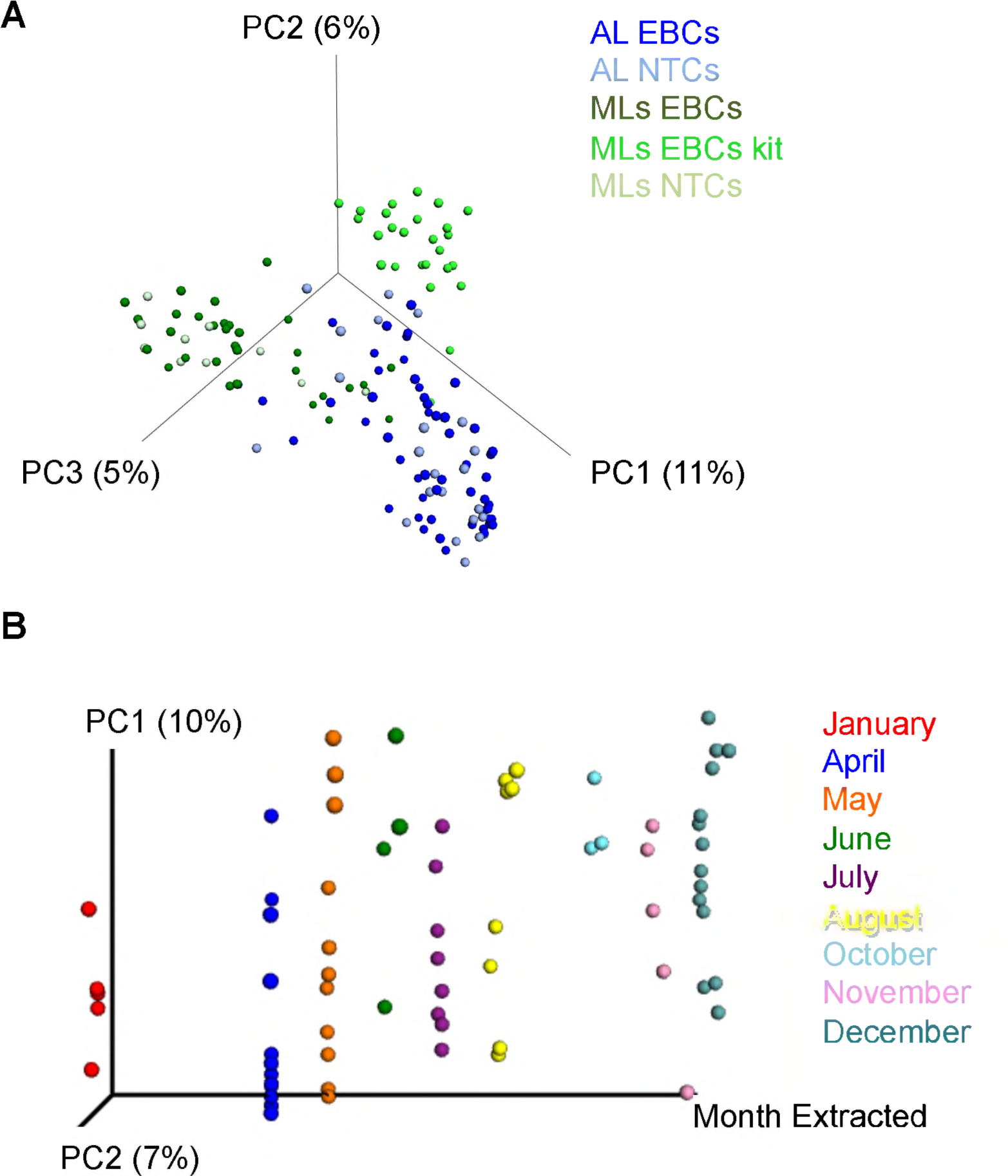
PCoA analysis of extraction method and seasonal variation on contaminant communities. The modern and ancient calculus samples were removed from the analysis presenting in Figure 3, and a PCoA plot was constructed of only control samples to identify differences between the extraction method and laboratory in control samples (A). (B) UniFrac values from controls samples (EBCs and NTCs) from the ancient laboratory over a five-year period (2012 – 2016) are colored on a PCoA plot according to month.

**Table 2:** Extraction methods contain unique taxa. OTUs identified as statistically significant (Kruskal-Wallis Bonferroni Corrected p- value <0.05) between the two extraction methods in the modern laboratory are listed. OTUs highlighted in green were significantly within the QG method, while highlighted OTUs were significant in the kit extraction method.

| OTU Taxonomy | Mean Seqs/Sample |  |
| --- | --- | --- |
|  | Kit | QG |
| Bacteria;p_Firmicutes;c_Bacilli;o_Bacillales | 6.535714 | 0.008696 |
| Bacteria;p_Firmicutes;c_Bacilli;o_Bacillales;f_Alicyclobacillaceae;g_Alicyclobacillus;s_ | 2.821429 | 0 |
| Bacteria;p_Firmicutes;c_Bacilli;o_Bacillales;f_Alicyclobacillaceae;g_Alicyclobacillus;s_ | 7.214286 | 0 |
| Bacteria;p_Firmicutes;c_Bacilli;o_Bacillales;f_Alicyclobacillaceae;g_Alicyclobacillus;s_ | 1.928571 | 0 |
| Bacteria;p_Firmicutes;c_Bacilli;o_Bacillales;f_Alicyclobacillaceae;g_Alicyclobacillus;s_ | 2.071429 | 0 |
| Bacteria;p_Firmicutes;c_Bacilli;o_Bacillales;f_Alicyclobacillaceae;g_Alicyclobacillus;s_ | 1.714286 | 0 |
| Bacteria;p_Proteobacteria;c_Gammaproteobacteria;o_Oceanospirillales;f_Halomonadaceae;g_Halomonas;s_ | 0.035714 | 0 |
| Bacteria;p_Actinobacteria;c_Actinobacteria;o_Actinomycetales;f_Pseudonocardiaceae;g_Pseudonocardia;s_ | 0.035714 | 0 |
| Bacteria;p_Proteobacteria;c_Gammaproteobacteria;o_Pseudomonadales;f_Moraxellaceae;g_Acinetobacter;s_ | 385.2857 | 0.026087 |
| Bacteria;p_Proteobacteria;c_Alphaproteobacteria;o_Rhizobiales;f_Bradyrhizobiaceae;g_s_ | 93.42857 | 0 |
| Bacteria;p_Proteobacteria;c_Alphaproteobacteria;o_Rhizobiales;f_Bradyrhizobiaceae;g_s_ | 91.85714 | 11.72174 |
| Bacteria;p_Firmicutes;c_Bacilli;o_Bacillales;f_Alicyclobacillaceae;g_Alicyclobacillus;s_ | 0.857143 | 0 |
| Bacteria;p_Firmicutes;c_Bacilli;o_Bacillales;f_Alicyclobacillaceae;g_Alicyclobacillus;s_ | 0.857143 | 0 |
| Bacteria;p_Proteobacteria;c_Betaproteobacteria;o_Neisseriales;f_Neisseriaceae;g_Vogesella;s_ | 184.4643 | 0 |
| Bacteria;p_Proteobacteria;c_Alphaproteobacteria;o_Rhizobiales;f_Bradyrhizobiaceae | 2.357143 | 0.008696 |
| Bacteria;p_Proteobacteria;c_Gammaproteobacteria;o_Pseudomonadales;f_Moraxellaceae;g_Acinetobacter;s_ | 395.3929 | 0.626087 |
| Bacteria;p_Proteobacteria;c_Alphaproteobacteria;o_Sphingomonadales;f_Sphingomonadaceae;g_Sphingobium;s_ | 160.75 | 33.24348 |
| Bacteria;p_Firmicutes;c_Erysipelotrichi;o_Erysipelotrichales;f_Erysipelotrichaceae;g_Allobaculum;s_ | 0.607143 | 0 |
| Bacteria;p_Firmicutes;c_Bacilli;o_Bacillales;f_Alicyclobacillaceae;g_Alicyclobacillus;s_ | 0.428571 | 0 |
| Bacteria;p_Verrucomicrobia;c_Verrucomicrobiae;o_Verrucomicrobiales;f_Verrucomicrobiaceae;g_Akkermansia;s_muciniphila | 0.5 | 0 |
| Bacteria;p_Proteobacteria;c_Gammaproteobacteria;o_Pseudomonadales;f_Moraxellaceae;g_Acinetobacter;s_ | 82.14286 | 0.73913 |
| Bacteria;p_Proteobacteria;c_Alphaproteobacteria;o_Rhizobiales;f_Bradyrhizobiaceae | 1.964286 | 1.721739 |
| Bacteria;p_Firmicutes;c_Erysipelotrichi;o_Erysipelotrichales;f_Erysipelotrichaceae;g_Allobaculum;s_ | 0.464286 | 0 |
| Bacteria;p_Firmicutes;c_Bacilli;o_Bacillales;f_Alicyclobacillaceae;g_Alicyclobacillus;s_ | 0.392857 | 0 |
| Bacteria;p_Cyanobacteria;c_Chloroplast;o_Streptophyta;f_g_s_ | 2 | 0.008696 |
| Bacteria;p_Cyanobacteria;c_Chloroplast;o_Streptophyta;f_g_s_ | 26.10714 | 1.547826 |
| Bacteria;p_Firmicutes;c_Bacilli;o_Lactobacillales | 1.964286 | 1.991304 |
| Bacteria;p_Proteobacteria;c_Gammaproteobacteria;o_Enterobacteriales;f_Enterobacteriaceae;g_s_ | 5.392857 | 23.46087 |
| Bacteria;p_Proteobacteria;c_Gammaproteobacteria;o_Pseudomonadales;f_Pseudomonadaceae;g_Pseudomonas;s_ | 0 | 0.008696 |
| Bacteria;p_Proteobacteria;c_Betaproteobacteria;o_Burkholderiales;f_Comamonadaceae;g_s_ | 0 | 0.008696 |
| Bacteria;p_Proteobacteria;c_Gammaproteobacteria;o_Pseudomonadales;f_Pseudomonadaceae;g_Pseudomonas;s_ | 0 | 0.008696 |
| Bacteria;p_Proteobacteria;c_Betaproteobacteria;o_Burkholderiales;f_Comamonadaceae;g_s_ | 0 | 0.008696 |
| Bacteria;p_Proteobacteria;c_Gammaproteobacteria;o_Enterobacteriales;f_Enterobacteriaceae;g_s_ | 5.571429 | 44.6 |
| Bacteria;p_Proteobacteria;c_Gammaproteobacteria;o_Oceanospirillales;f_Alcanivoracaceae;g_Alcanivorax;s_ | 0 | 0.017391 |
| Bacteria;p_Proteobacteria;c_Gammaproteobacteria;o_Enterobacteriales;f_Enterobacteriaceae;g_s_ | 4.785714 | 20.73043 |
| Bacteria;p_Proteobacteria;c_Gammaproteobacteria;o_Enterobacteriales;f_Enterobacteriaceae;g_s_ | 4.928571 | 23.37391 |
| Bacteria;p_Proteobacteria;c_Epsilonproteobacteria;o_Campylobacteriales;f_Helicobacteraceae;g_s_ | 0 | 0.017391 |
| Bacteria;p_Proteobacteria;c_Gammaproteobacteria;o_Enterobacteriales;f_Enterobacteriaceae;g_s_ | 0 | 0.017391 |
| Bacteria;p_Proteobacteria;c_Gammaproteobacteria;o_Oceanospirillales;f_Halomonadaceae;g_Halomonas;s_ | 0 | 0.026087 |
| Bacteria;p_Proteobacteria;c_Gammaproteobacteria;o_Enterobacteriales;f_Enterobacteriaceae;g_s_ | 1.892857 | 4.33913 |
| Bacteria;p_Proteobacteria;c_Betaproteobacteria;o_Burkholderiales;f_Oxalobacteraceae;g_Cupriavidus;s_ | 0 | 0.034783 |
| Bacteria;p_Proteobacteria;c_Gammaproteobacteria;o_Enterobacteriales;f_Enterobacteriaceae;g_s_ | 1.107143 | 5.921739 |
| Bacteria;p_Proteobacteria;c_Alphaproteobacteria;o_Rhodobacterales;f_Rhodobacteraceae;g_s_ | 0 | 0.026087 |
| Bacteria;p_Proteobacteria;c_Gammaproteobacteria;o_Alteromonadales;f_Alteromonadaceae;g_Marinobacter;s_ | 0 | 0.034783 |

#### Contaminant taxa change over time

Much of the variation identified in this study is laboratory-specific, so in order to test how seasonal changes, different researchers, or time might alter the microbial diversity observed in controls, we assessed the EBC and NTC records from the ancient lab facility over five years (2012-2016). Bacterial community structure in the ancient lab was linked to the researcher (adonis; p=0.001,R^2^=0.073), the extraction year (adonis; p=<0.01,R^2^=0.022), the extraction month (adonis; p=<0.001,R^2^=0.044; Figure 4B), and wet / dry seasons (adonis; p=0.001,R^2^=0.081). However, each of these signals was less significant and drove less variation within the data set when compared to the differences observed between laboratory facilities or between extraction methods. Very few specific taxa were significantly associated with temporal variation, although linked changes in overall diversity were observed. 32 OTUs were associated with the month in which the extraction was performed and were largely present during dry months (Oct-January; dominated by *Comamonadaceae* (2), *Bradyrhizobiaceae*(11), and *Gemmatimonadetes*(2) taxa; Kruskal-Wallis; Bonferroni corrected p=<0.05), while only two OTUs (*Thermobispora*and *Actinomycetales*taxa) were linked to wet seasons. Interestingly, five OTUs (*Leptotrichia*, *Comamonadaceae* (3), and *Burkholderia*) were also associated with the lab researcher (Kruskal-Wallis; Bonferroni corrected p=<0.05). While we cannot rule out the confounding nature of these variables (*e.g.*links between different researchers being more active in the laboratories at different times), these observations suggest that contaminant taxa change over time and need to be continually monitored, even in the cleanest molecular facilities.

### Discussion

#### Overview

While several studies have now reported on contaminant DNA within laboratory reagents, the systematic inclusion of extraction blank controls has not yet been widely embraced in metagenomic research. Several studies on human microbiota have been criticised for their lack of careful controls [14, 24, 27], as the unfounded results of such studies have potentially serious repercussions and have hindered scientific progress. A similar phenomenon occurred with the new field of ancient DNA in the early 1990s, when research teams, reviewers, and editors failed to adequately test for contamination [28–30], leading to many spurious results. This seriously undermined the credibility of ancient DNA research [23] and resulted in the formation of a robust set of guidelines [9]. Here, we surveyed the largest collection of extraction blank and no-template amplification negative control samples to date (n=144) with the goal of better describing contaminant DNA in microbiome studies to avoid pitfalls similar to those observed in the ancient DNA field.

#### Contaminant diversity remains underestimated

We identified 861 contaminant taxa over five years within a single ultra-clean laboratory facility. Before this publication, the largest collection of contaminant taxa was published by Salter *et al.*and included 93 contaminant genera [3]. Within our study, we found 71 of the taxa identified by Salter *et al.* across all labs and methodologies. However, only 29.5% of the Salter *et al.* taxa (21 of their 71 taxa) were identified as dominant taxa within our study across all methods and labs. This indicates that laboratory microbial contamination is not yet well described and is likely to be unique across different laboratories, protocols, seasons, and researchers. Of the 21 taxa shared across studies, four genera (*Ralstonia*, *Acinetobacter*, *Pseudomonas*, and *Stenotrophomonas*) have now been routinely identified in at least four of the six publications that examine laboratory contamination [2–4, 13, 31, 32]. All of these taxa are classified as Proteobacteria, as are 55% of the dominant contaminant taxa (38/69) identified within our study and 63% (34/92) within the Salter *et al.* study. While contamination is highly diverse, this finding indicates that Proteobacteria appear to be the most widespread source of laboratory contamination. Proteobacteria encompasses several families of bacteria that are known to be UV and oxidation resistant.

#### Analysing contaminants is critical for the successful interpretation of low-biomass samples

We identified several human oral microbiota taxa present in the commercial extraction kit, including *Fusobacterium, Streptococcus,* and *Corynebacterium* [33], while previous studies have previously identified additional human oral taxa contaminants, including *Haemophilus* and *Peptostreptococcus* [31]. Worryingly, one of these taxa in particular, *Fusobacterium*, has recently been identified both as a component of the ‘placental microbiome’, and as a component of breast cancer tissue, in low-biomass studies that did not consider background contamination from laboratory reagents or environments [22, 26, 34]. It remains unclear whether this taxon is a laboratory contaminant, or whether it can escape the oral cavity and contribute to inflammatory processes elsewhere in the body. Other non-oral taxa identified within this study as contaminants have also previously been reported as important taxa within studies that failed to use controls [35]. There is clearly a need for more detailed metagenomic studies, or the use of improved ‘oligotyping’ 16S rRNA gene analysis methods of contaminant taxa, to better identify specific strain differences and determine whether such taxa are contaminants or are actually present in the body and can cause systemic disease. The lack of contaminant assessment has already negatively impacted the metagenomics field [14], and it is critical that editors and reviewers are aware of this issue.

#### Bacterial DNA is still obtained from ultra-clean reagents in ultra-clean facilities – no facility is contaminant free

Contaminant taxa were identified in EBCs and NTCs within five different laboratory facilities, including a state-of-the-art, ultra-clean ancient DNA facility. In the latter, the specialized conditions and procedures did not prevent low levels of bacterial diversity, and a wide-range of contaminant taxa was still observed – with the dominant taxa all known to resist disinfectant measures, including treatment with aromatic or oxidative compounds (*i.e.*bleach) (*Acinetobacter* [36], *Comamonas* [37], or other disinfectant compounds (*Pseudomonas* [38])). These mechanisms of disinfection resistance have contributed to nosocomial infections in hospitals (*i.e. Acinetobacter* [39]) and to contamination of cell culture reagents (*e.g. Achromobacter* [40]). Of note, *Deinococcus*, a taxa that can notoriously survive UV irradiation [41], *Alicyclobacillus*, known to survive pasteurization [25], and other species known to degrade oxidative compounds (*e.g. Pasteurella* [42]) were not observed in the specialised ancient DNA facility, but were identified within the modern laboratory. While measures to reduce contamination have prevented the introduction of human-associated microorganisms into the ancient lab EBCs, these numerous strategies did not eliminate or completely prevent the introduction of bacterial contaminant DNA. This suggests that each research facility will likely contain unique microorganisms able to resist decontamination measures, although it is plausible that contaminant DNA could be routinely introduced into the facility from other source and represents living species found elsewhere, rather than in the actual facilities utilized in this study. Regardless, this finding reiterates that every laboratory is susceptible to bacterial DNA contamination and that researchers should consistently monitor the contamination present within their own facility as a best practice.

#### Non-kit approaches provide unique contaminant signals

In this study, we identified several taxa in a commonly used DNA extraction kit that were absent in the homemade ancient DNA extraction method. The ancient DNA method was developed to obtain more DNA from samples with low-endogenous DNA, and this and other similar extraction methods are now routinely applied in ancient DNA studies to examine ancient microbiota and metagenomes [11, 43, 44]. In this study, the ancient DNA method produced extraction blanks that had lower microbial diversity and were less likely to contain human oral taxa than extraction blanks generated using a commercial kit. This suggests that commercially available kits may contain more DNA contamination than homemade methods that source clean materials. It is likely that the assembly of kit-based reagents in a separate facility provides an additional opportunity to contaminate reagents with laboratory DNA. This also suggests that ancient DNA extraction methods and strategies could be applied in modern low-biomass studies to potentially reduce contaminants that originate from humans.

In the future, studies of low-biomass or low endogenous count routinely employ shotgun sequencing to better identify contaminant taxa, as strain-level identifications increase specificity in tracking contaminants. In many cases, the ancient DNA field has now shifted to utilizing shotgun DNA sequencing as the gold-standard method (12). Shotgun sequencing also produces many other important molecular signals (*e.g.* signatures of ancient DNA damage), functional analysis, and strain markers to delineate which species are endogenous and which are contaminants. For example, distinct strains within a single genus could be identified as either a contaminant or an endogenous species, which would be critical for examining oral species in low-biomass tissues. In addition, damage profiles of DNA contamination could be used to distinguish fragmented, extracellular DNA within reagents versus species living within the laboratory. Current approaches aimed at eliminating contamination in shotgun sequenced metagenomes have had varied levels of success (reviewed in [3]), and new bioinformatic tools and models will undoubtedly improve our ability to identify and account for contaminant signals within metagenomic data sets (45). However, the need to routinely include EBCs and NTCs within microbiome data sets will likely always be necessary when examining low biomass samples, even when other methodologies, such as shotgun metagenomic sequencing, are applied.

#### Contamination assessment needs to be routinely reported as a publication requirement

Contaminant sequences introduced during sample processing and library construction significantly contribute to signals from biological samples, especially those that are low-endogenous or low-biomass in nature. This study confirms that contaminant taxa that are unique to the extraction method and facility, are related to the material being extracted, and change over time within a single facility, although these levels of contamination can be somewhat mitigated by routine decontamination measures of the facility and potentially the reagents themselves (46). Therefore, the presence of contaminants needs to be considered in all future studies of both human and environmental microbiota. We recommend that all researchers routinely record potential sources of contamination DNA (reagent batches or lot numbers; dates of extractions and amplifications; researchers performing such duties, *etc.*) and critically propose that researchers routinely include extraction blank controls during the extraction process to monitor the bacterial DNA introduced into their samples. Minimally, one control should be included in at least every batch of extractions and amplifications performed. Adding carrier DNA into control samples may also improve contaminant DNA detection (47). If controls were not included in existing data sets, an assessment of previously identified contaminant taxa within study datasets should also be minimally included in the published analysis. For example, researchers could report how many known contaminant taxa are present within a dataset or provide evidence to demonstrate that the removal of known contaminants does not impact the sample signal or conclusions of the paper. To facilitate this process, we have included a text file that includes a list of all the contaminant taxa observed here, as well as a separate file of only the dominant taxa. The inclusion of negative extraction blank controls should be regarded as minimal requirements for any metagenomics research and should become standard requirements of reviewers and journal editors.

## Acknowledgements

We would like to acknowledge Paul Gooding at the Australian Genomic Research Facility for technical help during DNA sequencing. This research was funded by the Australian Research Council (L.S.W; A.C.).

## Data Accessibility

QIIME demultiplexed sequences (16SContam_seqs_forpub.fna), a phylogenetic tree of representative sequences (rep_set.tre), a biom table (otu_table_clean.biom), and sample metadata (SampleInformation_20180820.txt) can be accessed from https://figshare.com/account/articles/7283816 (doi: 10.25909/5bdaa4431a941).

## Author Contributions

LSW and AGF conceived of the study. LSW, AGF, RE, JY, CS, MHD, and CA contributed samples and completed lab work. LSW, LA, and JB completed bioinformatic analysis of the data. LSW wrote the paper, and all authors edited and contributed to the final manuscript.

